# Shifting to biology promotes highly efficient iron removal in groundwater filters

**DOI:** 10.1101/2024.02.14.580244

**Authors:** Simon Müller, Francesc Corbera-Rubio, Frank Schoonenberg-Kegel, Michele Laureni, Mark C.M. van Loosdrecht, Doris van Halem

**Affiliations:** Delft University of Technology, The Netherlands; Vitens N.V., The Netherlands

## Abstract

Rapid sand filters are established and widely applied technologies for groundwater treatment. Conventionally, intensive aeration is employed to provide oxygen for the oxidation and removal of the main groundwater contaminants. While effective, intensive aeration promotes flocculent iron removal, which results in iron flocs that rapidly clog the filter. In this study, we operated two parallel full-scale sand filters at different aeration intensities to resolve the relative contribution of homogeneous, heterogeneous and biological iron removal pathways, and identify their operational controls. Our results show that mild aeration in the LOW filter (5 mg/L O_2_pH 6.9) promoted biological iron removal and enabled iron oxidation at twice the rate compared to the intensively aerated HIGH filter (>10mg/L O_2,_pH 7.4). ESEM images showed distinctive twisted stalk-like Fe solids, biosignatures of *Gallionella ferruginea*, both in the LOW filter sand coatings as well as in its backwash solids. In accordance, 10 times higher DNA copy numbers of *G. ferruginea* were found in the LOW filter effluent. Clogging by biogenic FeOx was slower than by chemical FeOx flocs, resulting in lower backwash frequencies and yielding four times more water per run. Ultimately, our results reveal that biological Fe^2+^ oxidation can be actively controlled and favoured over competing physico-chemical routes. The resulting operational benefits are only starting to be appreciated, with the counterintuitive higher oxidation rates and water yields at lower aeration regimes, and the production of more compact and practically valuable FeOx solids being of outmost interest.

## 1. Introduction

Groundwater provides safe drinking water to more than half of the world’s population (Connor, 2015), but often it naturally contains dissolved iron (as Fe^2+^). Exposure to Fe via drinking water is not harmful, yet the WHO recommends Fe concentrations not to exceed 0.3 mg/L (World Health Organization, 2022). This is to avoid orange-brown discoloration of tap water and to prevent deposition of solids in distribution systems. Fe removal is achieved by oxidation of dissolved Fe^2+^ to insoluble Fe(III)(oxihydr)oxides (or FeOx), followed by separating these FeOx solids by sand filtration. When dissolved oxygen (DO) is the oxidant, three main Fe^2+^ oxidation pathways can be identified (Figure 1): (i) abiotic *homogenous* oxidation, (ii) *heterogeneous* oxidation, catalysed by existing FeOx surfaces, and/or (iii) *biological* oxidation by microorganisms (Van Beek et al., 2012). Where homogeneous oxidation occurs in the water phase, heterogeneous and biological oxidation occur on the surface of the filter sand grain. Each oxidation pathway produces morphologically different FeOx and can be found either in the (pore) water or on the sand grain. In full-scale filters, different Fe^2+^ oxidation pathways have been observed to co-occur.

**Figure 1.**
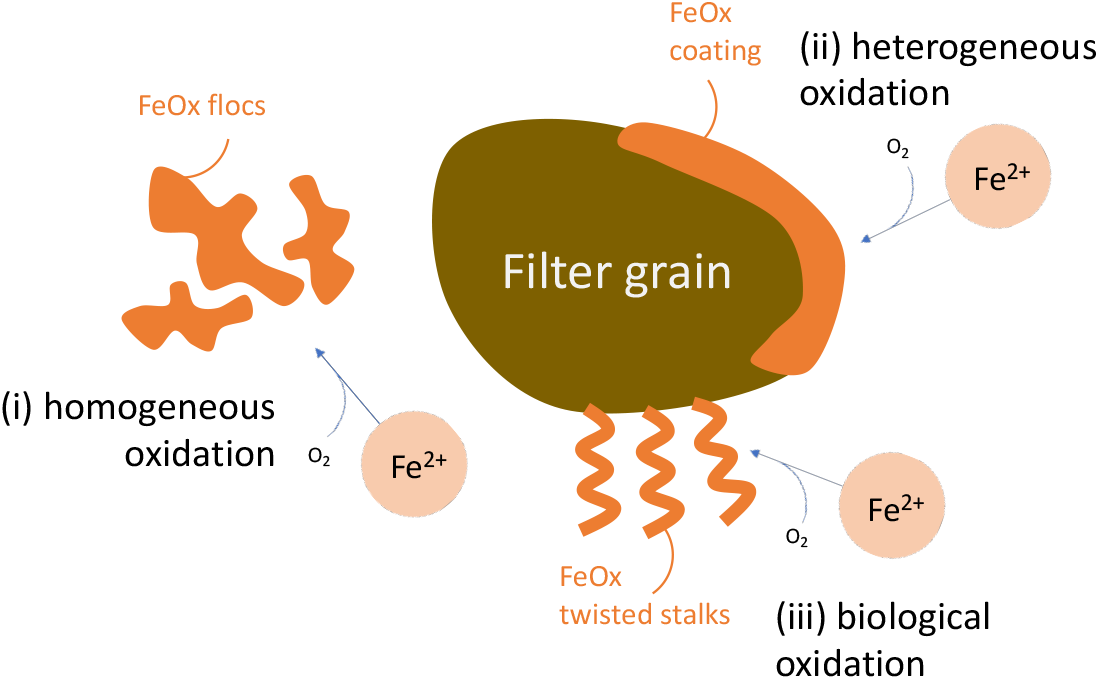
Three main Fe^2+^ oxidation pathways during aeration-filtration of groundwater: homogenous oxidation in the water, heterogeneous oxidation on FeOx, and biological oxidation by microorganisms.

The traditional view on Fe removal from groundwater is based on the first mechanism, homogeneous oxidation, which is favoured by intensive aeration prior to filtration. Aeration strips dissolved, undesired gases (CH_4_, H_2_S) and partially removes dissolved carbon dioxide (CO_2_), which raises the pH. Simultaneously, oxygen (O_2_) dissolves into the reduced groundwater. The changes in pH and reduction-oxidation-potential facilitate the oxidation of Fe^2+^ by dissolved oxygen (DO), either directly or indirectly, and thus produce FeOx (Appelo and Postma, 2005), which can eventually be retained in the filter. Fe flocs in a clogged bed are removed by periodic backwashing, which is achieved with a reversed flow of water at a high velocity or with water combined with air. Homogeneous Fe^2+^ oxidation creates watery Fe sludge as by-product, which needs additional treatment.

Fe removal on the surface of filter sand grains, either via FeOx or by microorganisms, has been proposed to result in less frequent backwashing and, higher filtration rates (Mouchet, 1992; Sharma, 2001). Although surface-related Fe removal has been applied in France (Mouchet, 1992), Belgium (Van Beek et al., 2016), Denmark (Søgaard et al., 2000) and other countries for many years, it is often treated as a single mechanism, termed either *biological* (Mouchet, 1992; Søgaard et al., 2000) or *adsorptive* (Sharma, 2001). Van Beek (2016) attempted to quantify the individual mechanisms and concluded that the contribution of biological removal was highest around pH 7.5, contradicting Mouchet’s (1992) observation that biological Fe removal usually requires a pH around 7 or below. De Vet et al. (2011b) investigated fully aerated trickling filters and found low groundwater temperatures to delay chemical oxidation, whereas biological Fe^2+^ oxidation by *Gallionella* spp. was promoted.

In groundwater filters the boundaries between abiotic and biological Fe removal are still not well understood, neither their potential positive or negative interactions (Emerson and De Vet, 2015). The contribution of different Fe oxidising bacteria to total FeOx production has been studied in the laboratory at very low DO concentrations (Druschel et al., 2008; Maisch et al., 2019). Biological Fe^2+^ oxidation seems to dominate under these conditions, but whether exclusive biologically-driven Fe removal is possible under conditions relevant for groundwater treatment remains unknown. We hypothesize that pH and oxygen concentration are the key parameters that dictate the dominant Fe oxidation mechanism in sand filtration. To this end, we designed and operated two full-scale sand filters, namely HIGH (DO<10 mg/L, pH 7.4) and LOW (DO<5 mg/L, pH 6.9) to untangle homogenous, heterogeneous and biological Fe removal pathways. Ultimately, the overarching goal of this study was to gain mechanistic knowledge on how to control and promote biological Fe^2+^ oxidation in sand filters, paving the way towards novel design and efficient operation of groundwater filters.

## 2. Materials and methods

### 2.1. Groundwater composition

The two full-scale groundwater sand filters investigated in this study were operated at a Dutch groundwater treatment plant to produce drinking water. Groundwater is abstracted from a well field tapping into an anaerobic aquifer, with both DO and NO_3-_ below detection limit. Groundwater pH at this location is on average 6.72, and Fe^2+^, NH^4+^ and Mn^2+^ are on average 25.4, 2.19 and 0.32 mg/L, respectively. Table 1 provides an overview of the average groundwater composition for this treatment plant.

**Table 1.** Average groundwater composition.

|  | pH | NH<sub>4</sub><br>mg N/L | NO<sub>3</sub><br>mg N/L | Fe<br>mg/L | Fe(II)<br>mg/L | Mn<br>mg/L | SO<sub>4</sub><br>mg/L | CH<sub>4</sub><br>mg/L | PO<sub>4</sub><br>mg P/L | Ca<br>mg/L | Si<br>mg/L | HCO<sub>3</sub><br>mg/L |
| --- | --- | --- | --- | --- | --- | --- | --- | --- | --- | --- | --- | --- |
| Average | 6.72 | 2.19 | < 0.2 | 26.7 | 25.4 | 0.32 | 5.5 | 14.1 | 0.32 | 73.3 | 9.9 | 287 |
| Std.dev.<sup>1</sup> | 0.06 | 0.05 | NA | 1.5 | 3.1 | 0.02 | 0 - 16 | 2.8 | 0.04 | 2.9 | 0.4 | 11 |
| N | 137 | 32 | 32 | 137 | 137 | 137 | 32 | 137 | 24 | 32 | 32 | 32 |

<sup>1</sup> range instead of standard deviation shown for SO<sub>4</sub>

### 2.2 Full-scale filters

We investigated two parallel full-scale filters with different aeration intensities (Figure 2). The reference filter (HIGH) received water after plate aeration (100 m^3^/h, 4 m^2^), whereas filter LOW received water after membrane degassing (two 3M Liqui-Cel EXF-14x40 Hollow Fibre Membrane Contactors; first stage N_2_, second stage N_2_/CO_2_ sweep gas) and mild cascade aeration on top of the filter. The aim of LOW was to supply stoichiometrically sufficient DO to oxidise the Fe^2+^ in the groundwater (5 mg/L). For HIGH, DO concentrations reached saturation (>10 mg/L). Filter HIGH consisted of a dual bed, with 0.75 m anthracite on top of 1.25 m sand. Filter LOW was build-up of a 2.35 m single bed of coarse sand. The average filtration rates were 5 m/h for the HIGH filter and 10 m/h for the LOW filter. Further details of filter design and operational conditions are provided in Table 2. Both filters were backwashed periodically, Figure S1 details the backwash programs. Backwash water is treated in sedimentation basins and lamella flocculation to meet discharge regulations. The effluent of both filters is mixed in a storage reservoir and subsequently treated by tower aeration, secondary filters (for NH_4+_ and Mn^2+^ removal) and granular activated carbon filtration for colour removal. The produced drinking water is distributed to consumers without further disinfection, as well as used for backwashing.

**Table 2.** Filter design and process conditions.

|  | HIGH | LOW |
| --- | --- | --- |
| Filter media |  |  |
| Material, diameter | Anthracite, 2.5 – 4 mm<br>Sand (quartz), 0.8 – 1.25 mm | Coarse sand (quartz), 1.7 – 2.5 mm |
| Height | 0.75 m<br>1.25 m | 2.35 m |
| Media age (in Jan 2021) | 1 year<br>2 years | 2 months |
| Supernatant height | 30 – 60 cm | 30 - 60 cm |
| Filter surface area | 5 $\text{m}^2$ | 5 $\text{m}^2$ |
| Filtration rate | 5 m/h | 10 m/h |

**Figure 2.**
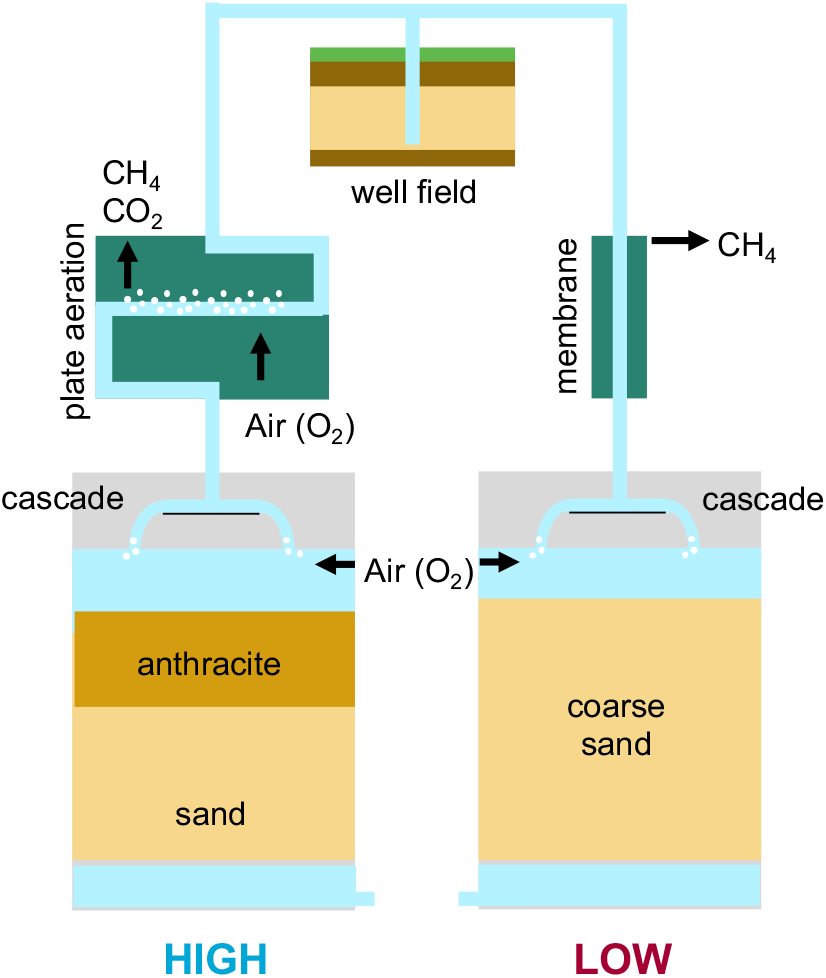
Overview of the two parallel full-scale treatment schemes investigated: The HIGH filter, a dual media filter (anthracite/sand) operated at approx. 5 m^3^/h/m^2^ with upstream plate aeration and the LOW filter, a single media filter (sand) operated at approx. 10 m^3^/h/m^2^.

### 2.3 Sampling procedures

Water samples were taken during normal operation, at the end of the filter run, i.e. prior to backwashing. Samples were collected from in-line sample points (raw water, after plate aeration, after membranes) as well as over the depth of the filter via sample points at the wall of the filter. The pH, temperature, redox potential and the DO concentration were measured on-site using a flow-cell attached to the sample point to avoid contact with the atmosphere. Samples for the analysis of ammonium (NH_4+_), nitrite (NO_2-_) and nitrate (NO_3-_) were filtered (0.2 µm polyethersulfone; MACHEREY-NAGEL, Germany) and stored on ice. Unfiltered 500 mL samples were collected for DNA extraction and stored on ice. Filtered and unfiltered samples for the analysis of metals and phosphate were taken and immediately acidified with ultrapure HNO_3_to a final concentration of 1% to avoid oxidation.

The flow was stopped at the end of the filter run, and filter medium was sampled using a disinfected stainless steel peat-sampler. The top 120 cm of each filter were sampled in 20 cm segments, transferred into polypropylene tubes and stored in a cooler. Samples for solid or DNA extraction were stored in RNase-free polypropylene tubes and stored on ice. The filters were subsequentially backwashed and the backwash suspension was sampled using two peristaltic pumps over the entire program. The tubing was positioned in the supernatant, halfway the radius of the filter. For the analysis of total Fe and Mn, averaged samples were taken every minute.

### 2.4 On-site Fe^2+^ oxidation experiments

For Fe^2+^ oxidation batch experiments, 1 L of aerated water was obtained after plate aeration (for HIGH) and filter supernatant (for LOW) to assess oxidation kinetics in the supernatant water. Experiments were executed immediately after sampling to minimize Fe^2+^ oxidation prior to the experiment. Glass beakers were wrapped in aluminium foil and placed on a magnetic stirrer at low speed open to the atmosphere. Fe^2+^ oxidation was monitored by taking regular samples for a period of 20 minutes. Dissolved oxygen, pH and temperature were monitored throughout the experiment. Experimental conditions for HIGH were DO 9 – 10.5 mg/L, pH 7 – 7.4 and temperature 11 – 15 °C, and for LOW the conditions were DO 4.5 – 5 mg/L, pH 6.7 – 7.1 and temperature 11 – 14 °C. Samples were immediately filtered over a 0.2 µm syringe filter. The results were simulated according to Stumm and Morgan (1976) at pH 7 – 7.4 and DO 10 mg/L for HIGH, and at pH 6.7 – 7.1 and DO 5 mg/L for LOW.

### 2.5 Fe^2+^ adsorption isotherms

Fe^2+^ adsorption isotherms were determined with fresh filter medium at room temperature at pH 7 (buffered with CO_2-HCO3_). For both types of filter media, duplicates of 1, 2, 4 and 8 g of wet grains were weighed into 100 mL serum bottles. Inside an anaerobic chamber, the bottles were filled with 100 mL buffer (5 mM HCO^3^) and Fe(II) stock (FeCl_2_ · 4 H2O) to a final concentration of 29 mg/L Fe^2+^. Clamped bottles were shaken for 24 h at 150 rpm. Bottles were opened in the anaerobic chamber and the Fe concentration in 0.2 µm filtered water samples was analysed colourimetrically (APHA, 2018). The remaining water was decanted and the dry weight of the solids was determined after 24 h at 95 °C.

### 2.6 Wet chemical analysis

The pH and temperature, the DO concentration, and redox potential were measured on site using SenTix® 980, FDO® 925 and a SenTix® ORP 900 (Pt & Ag/AgCl) probe, respectively (WTW, Germany). The concentrations of Fe, manganese (Mn), and phosphorus (P) were measured by ICP-MS (Vitens laboratory). The unfiltered samples for total metals and P were filtered after being stored for at least 2 days (acidified to 1% HNO_3_). Ammonia (NH_4+_), nitrite (NO_2-_) and nitrate (NO_3-_) concentrations were stored at 4 °C and analysed within 24 hours using photometric analysis (Gallery Discrete Analyzer, Thermo Fisher Scientific, USA).

### 2.5 Solid characterization

All samples were stored on ice during transport and transferred to a freezer or fridge within 12 hours. The filter medium samples for Mössbauer, XRD, and qPCR analysis were stored at -20 °C. The samples for solid extraction, Fe^2+^ adsorption, and microscopy were stored at 4 °C. For light microscopy, single grains were dried for 2 days at room temperature and several drops of the fresh suspension were dried on glass slides at room temperature. Subsamples of 30 mL bulk filter medium were freeze dried for composition analysis. Backwash suspension samples were stored at 4 °C and processed within 2 days. The samples were resuspended and known volumes were filtered over 0.45 µm cellulose acetate membrane filters (Whatman™, UK). The solids on the filters were stored at -20 °C, freeze dried, and transferred into an anaerobic chamber (95% Ar/5% H_2_ with Pd catalyst; COY, USA).

Coatings and dried backwash solids were extracted using citrate-buffered dithionite following the procedure of (Claff et al., 2010). The elemental composition was determined by ICP-OES. For total Fe/Mn in backwash suspensions, the slurries were dissolved in aqua regia for 2 h at 98°C and analysed by ICP-MS.

Light microscopy images were taken using a VHX-5000 digital microscope (Keyence) with an VH-Z20UR lens (images of grains) or VH-Z100UR lens (images of flocs or stalks). Environmental SEM (ESEM) images were taken with a FEI Quanta FEG 650 SEM at 0.5 °C and 6 – 8 mbar H_2_O atmosphere.

For Mössbauer spectroscopy and X-ray diffraction (XRD), filter media were thawed, dried and processed inside an anaerobic chamber. The coating was removed from the filter medium by gentle grinding with a mortar and pestle. The quartz or anthracite grains were separated manually and the solids were powdered carefully. For Mössbauer, samples were diluted with powdered sugar (target 80 mg Fe) and sealed with Kapton tape in a Cu ring (20 x 2 mm). Transmission ^57^Fe Mössbauer spectra were collected at room temperature with a conventional constant-acceleration spectrometer using a 57Co(Rh) source. Velocity calibration was performed using an α-Fe foil. The Mössbauer spectra were fitted using the Mosswinn 4.0 program. For XRD, powdered samples were stored in gas tight tubes. For the measurement, the powders were deposited on a Si510 wafer and analyzed with a Bruker D8 Advance diffractometer using a rotating sample stage. Diffraction patterns were collected from 5 – 130°, step size 0.030 ° 2θ, counting time per step 2 s.

### 2.6 DNA extraction and qPCR

For water samples, 500 mL was filtered over a 0.22 µm polycarbonate membrane filter (Millipore). In case of visible Fe deposits, 10-20mL ammonium oxalate was added for 10 s, prior to flushing with 10mL DNA/RNA free water (Invitrogen). This was repeated in case Fe deposits were still visible. For the filter grain samples, 1 g was added to 10 mL of DNA/RNA free water and placed with beads in a sonification bath. After sonication for 2 min, the released bacteria, which were attached to the particles, entered the solution. The solution was subsequently filtered and treated with oxalate, like the water samples. Power Biofilm DNA Isolation kits (Qiagen) were used for DNA isolation and DNA samples were stored in -80°C freezer. Gal16sQr and Gal16sQf primers, and Gal16sQ probe (Biolegio) were used for determination of *Gallionella ferruginea* with qRealTime PCR (CFX). Amplification was performed by initial denaturation for 3 min at 95 °C, followed by 43 cycles of amplification (10 s denaturation at 95 °C; 60 s annealing at 60 °C; 20 s elongation at 72 °C). To quantify the number of *G. ferruginea* in a sample, a standard dilution series with a known amount of *G. ferruginea* of the same target is used, which is analyzed by PCR in duplicate simultaneously with the samples. The CFX Maestro software (Biorad) calculated, by using the Cq values of the standard, the amount of *G. ferruginea* DNA copies in a sample.

## 3 Results

### 3.1 Higher flows and longer runtimes for LOW

Two full-scale filters were operated for over a year treating anaerobic groundwater. Initially, both filters operated at a filtration rate of 5 m/h, but after 2 months the filtration rate of LOW was increased to 10 m/h (Figure 3A). The volume produced per run varied per backwash cycle (Figure 3B)was typically in the range 540-700 m^3^ for HIGH, while for LOW this varied in the range of 1800-3500 m^3^, depending on the run time and filtration rate chosen. The backwash frequency depended on the build-up of hydraulic resistance due to clogging; backwash was initiated when supernatant height reached 60 cm. These operational parameters show that the LOW filter could be operated at a higher flowrate (10 vs 5 m/h) and with double to triple runtimes between backwashes, producing up to 4 times more water per run than the HIGH filter.

**Figure 3.**
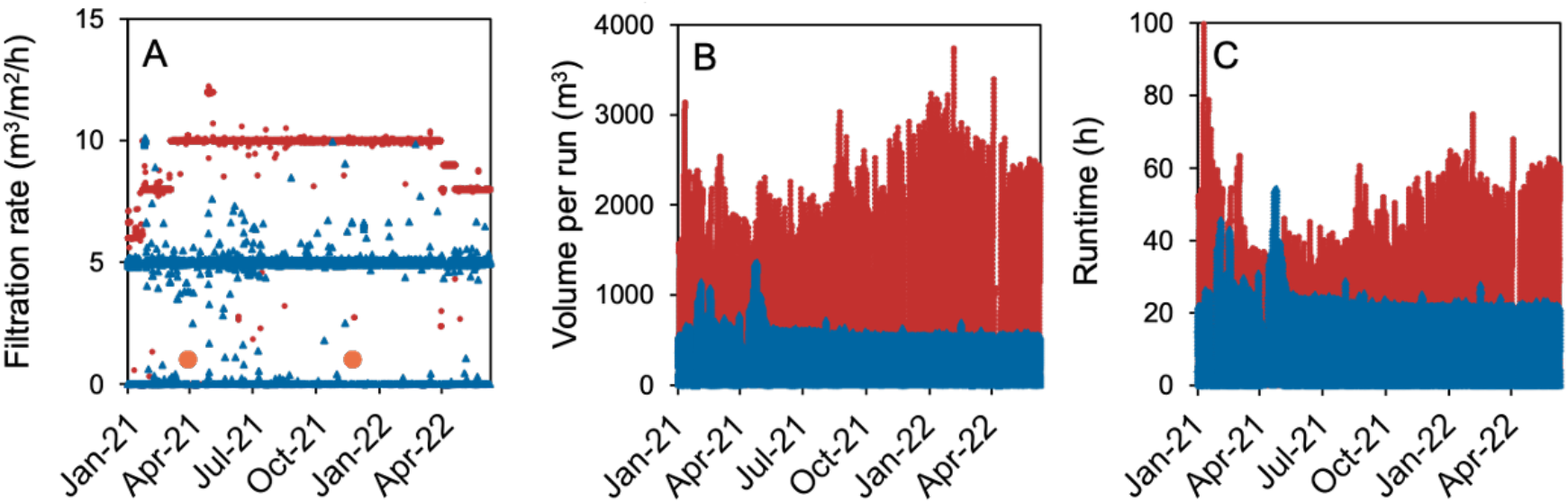
Operational parameters of LOW (red) and HIGH (blue) filters over time: (A) filtration rate (volumetric load) in m^3^ groundwater per m^2^ filter surface per hour, (B) volume of water filtered until backwashing, and (C) time until backwashing (runtime). The two sampling days in the presented study are indicated by orange circles.

### 3.2 Lower pH and dissolved oxygen in LOW filter

Figure 4 shows the depth profiles of oxygen, pH and redox potential in the raw water, after gas exchange and along the filter depth. The raw groundwater is anaerobic (DO and NO_3−_ below detection limit, 14 mg/L CH_4_) and slightly acidic (pH 6.7). Plate aeration prior to the HIGH filter increased the pH to 7.4 due to partial CO_2_ removal. DO concentration simultaneously increased to near saturation (>10 mg/L). Membrane degassing of CH_4_ prior to the LOW filter only had a minor effect on the pH and did not introduce DO. No changes in redox potential were observed during gas exchange. Once the water was mildly aerated on top of the LOW filter, DO increased to 5 mg/L and pH reached its maximum at 6.9. In both filters, pH decreased in the first meter of filter bed. Beyond this depth, pH stabilised, yielding a filtrate pH of 6.9 for HIGH and 6.7 for LOW. The DO concentrations dropped to < 1 mg/L in the top meter of both filters. A difference perceived between the two filters was that DO did not fall below 0.5 mg/L in LOW but was close to 0 mg/L in the effluent of HIGH. The redox potential increased in similar trends over depth in both filters, reaching close to 300 mV (vs SHE) in the effluent.

**Figure 4.**
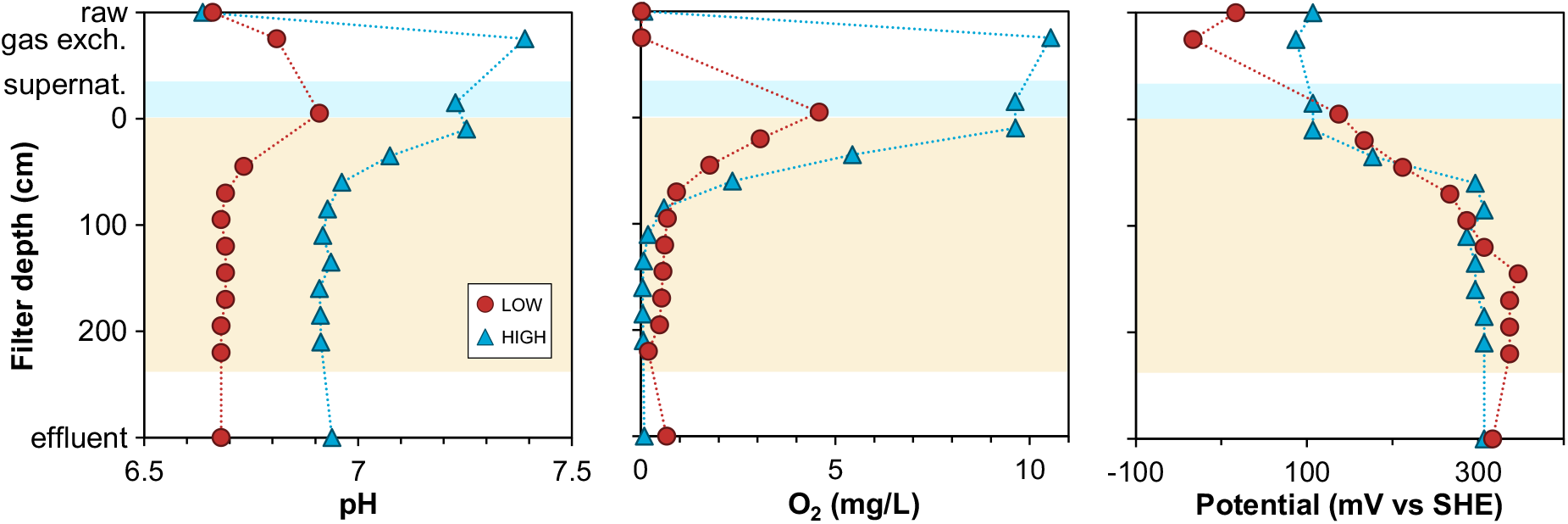
Profiles of measured water quality parameters in raw water, after gas exchange, and over the depth of both filters (LOW vs. HIGH). Subplots show the pH, the dissolved oxygen concentration (O2) and the redox potential. Water temperature was 10.7 – 11.3 °C.

### 3.3 Particulate Fe in water of HIGH filter

Groundwater Fe^2+^, at concentrations up to 30 mg/L, was removed in the top 60 cm of both filters (Figure 5), even though a double load was applied in the LOW filter compared to the HIGH filter. Depth profiles for dissolved and particulate Fe looked distinctly different for both filters. In the HIGH filter, a shift was observed from dissolved Fe in the raw water to flocculent, particulate Fe in the supernatant and in the top of the filter bed. This removal process is ascribed to homogeneous Fe oxidation and precipitation, known to occur at pH>7 and at high DO concentrations (Van Beek et al., 2016). On the contrary, dissolved iron removal in LOW was not accompanied by the formation of particulate Fe. The absence of formation of particulates in the water phase indicates that surface-related oxidation pathways dominate in filter LOW, *i*.*e*., heterogeneous and/or biological Fe^2+^ oxidation.

**Figure 5.**
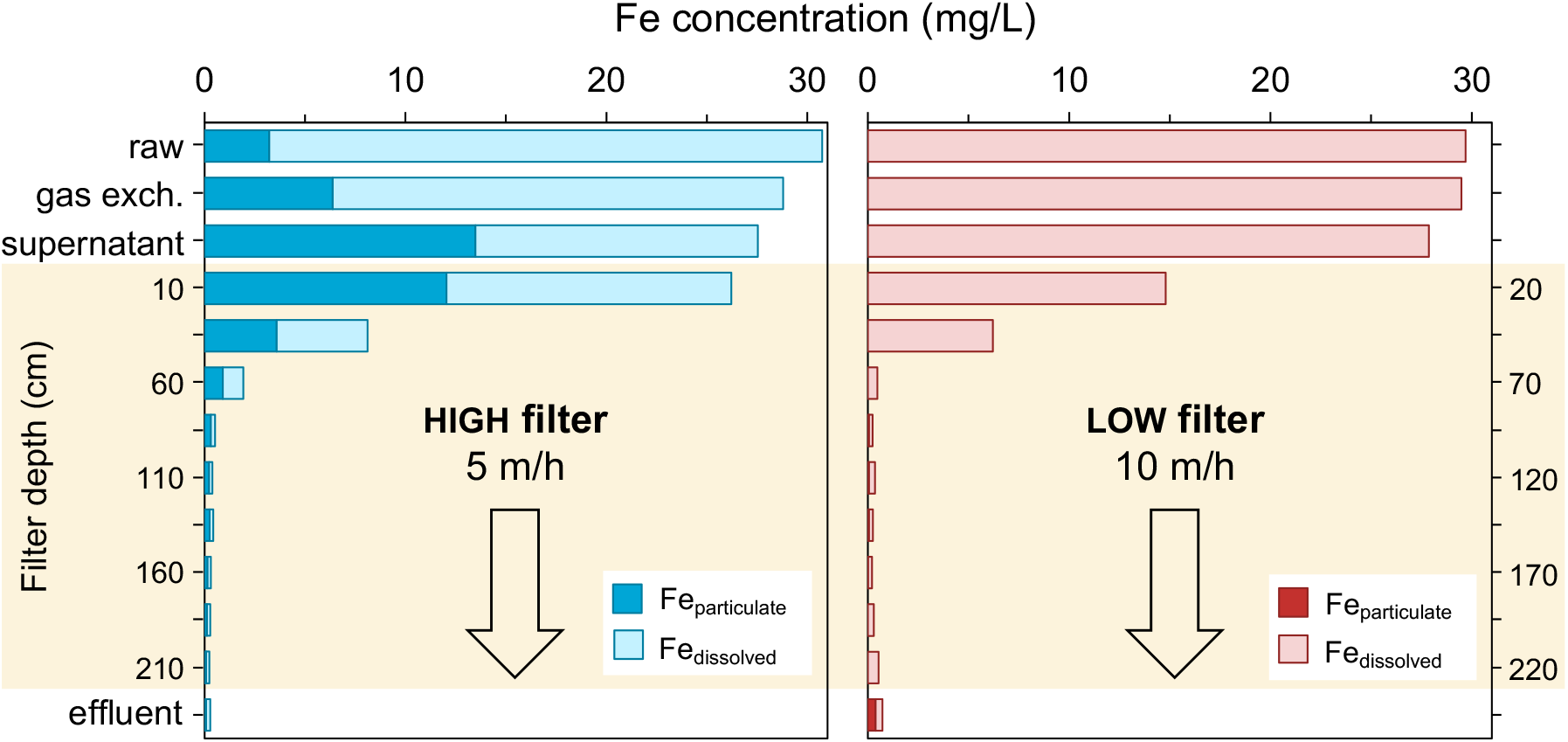
Fe concentrations (x-axis) measured along the parallel treatment schemes HIGH filter and LOW filter (y-axis). Dark bars indicate particulate Fe (>0.2 µm), light bars indicate dissolved Fe (<0.2 µm). ‘raw’ indicates raw water concentrations, ‘gas exch.’ the main gas exchange steps of plate aeration (HIGH) and membrane degassing (LOW).

### 3.4 Homogeneous oxidation in supernatant of filter HIGH

Fe^2+^ oxidation in the supernatant of both filters was simulated in batch experiments (Figure 6). Hardly any Fe^2+^ oxidation was found for LOW within the residence time in the supernatant (approx. 1-3 min). For filter HIGH, the water had a longer retention time prior to filtration, due to a lower flow rate and longer pathway from gas exchange to filter (approx. 6-10 min). In addition to residence time, pH also plays a major role in the homogeneous Fe^2+^ oxidation rate, as illustrated by the in-silico simulations for LOW (pH 6.7 and 7.1), and HIGH (pH 7.0 and 7.4). The lower pH and DO in the LOW filter yielded considerably slower homogeneous Fe oxidation, explaining the absence of particulate Fe in the LOW filter. This confirms that while in HIGH homogenous oxidation dominates, the fast Fe removal observed in the LOW filter was achieved by mechanisms other than homogeneous oxidation.

**Figure 6.**
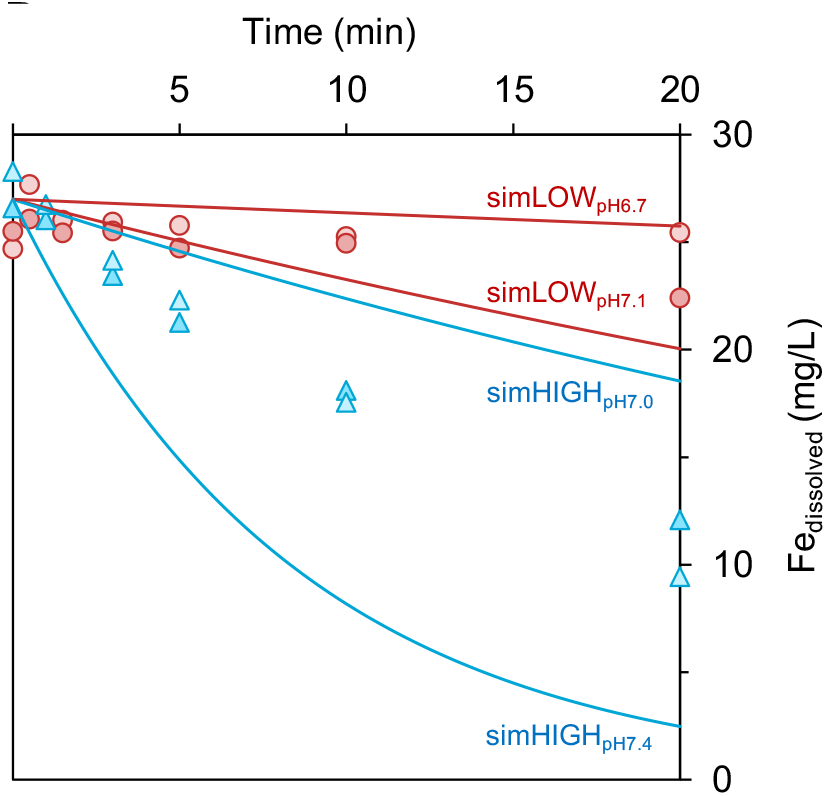
Dissolved Fe (Fe < 0.2 µm) (symbols) over time in on-site batch experiments with plate aeration effluent (HIGH; blue marks) and filter supernatant (LOW; red marks). Experimental conditions for HIGH: O_2,_ 9 – 10.5 mg/L; pH, 7 – 7.4; Temp., 11 – 15 °C, for LOW: O_2,_ 4.5 – 5 mg/L; pH, 6.7 – 7.1; Temp., 11 – 14 °C. Solid lines indicate the modelled Fe^2+^ concentrations (HIGH: pH, 7.0 – 7.4; O_2_, 10 mg/L; LOW: pH, 6.7 – 7.1; O_2_, 5 mg/L).

### 3.5 Dissolved phosphate enters LOW filter

In the HIGH filter, dissolved phosphate (PO_43-_), measured as total dissolved P, was completely removed from the water phase directly after plate aeration (354 µg/L; Figure 7A). Hence, no dissolved P entered the filter bed. Instead, P adsorbed to particulate Fe, as reflected in the increase in particulate P after gas exchange (Figure 7B). As a result, P removal in HIGH follows the removal of particulate Fe over the height of the filter bed. On the contrary, only a fraction of dissolved P disappeared (45 µg/L) prior to filtration in LOW, while the majority was removed within the filter bed. The removal profile of dissolved P in LOW resembled the removal profile for dissolved Fe, with complete removal at 80 cm depth. In both filters the removal of P is clearly linked to Fe removal, and consequently a shift in Fe oxidation pathway affects P removal.

**Figure 7.**
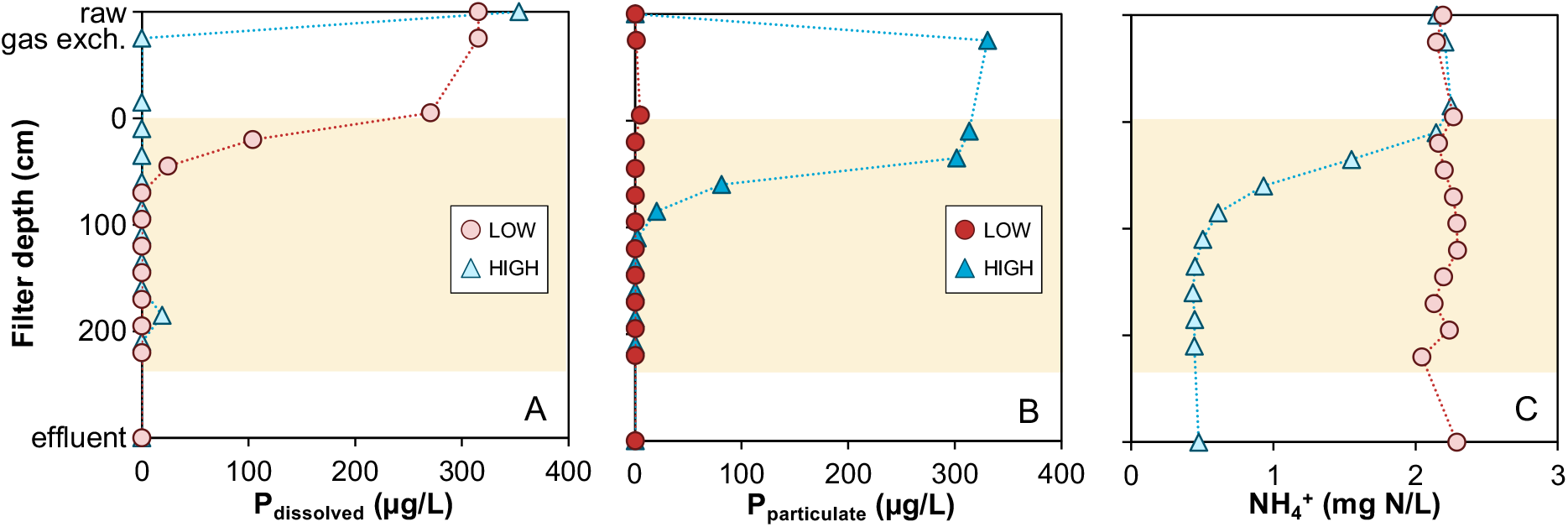
Concentration profiles (x-axis) of (A) dissolved, (B) particulate phosphorus (as total P) and (C) ammonium (NH_4_ ^+^) along the parallel treatment schemes HIGH filter and LOW filter (y-axis). Dark bars indicate particulate phosphate (>0.2 µm), light bars indicate dissolved phosphate (<0.2 µm). ‘raw’ indicates raw water concentrations, ‘gas exch.’ the main gas exchange steps of plate aeration (HIGH) and membrane degassing (LOW).

### 3.6 No nitrification in LOW filter

Figure 7C depicts the removal of NH_4+_ for the two filters. Prior to filtration, no nitrification was observed and 2.4 mg/L N-NH_4_/L entered both filter beds. In filter HIGH, over 75% N-NH_4_/L was removed in the first 120 cm of bed, while no NH_4+_ oxidation occurred in the LOW filter. As a consequence of aerobic NH_4+_ oxidation in HIGH, NO_2-_ (5-57 µgN/L) and NO_3-_ (0.5-2.1 mgN/L) were also detected across the filter depth. The detection of NO_2-_ in the effluent of HIGH indicates that nitrification was not complete, likely due to oxygen limitation. In addition to NH_4+_, dissolved manganese was present in the raw water. In both filters minor Mn^2+^ removal was observed only (30-70µg/L). Any residual NH_4+_, NO_2-_ or Mn^2+^ in the effluent of both primary filters was removed in the secondary aeration – filtration step (not the scope of this study).

### 3.7 LOW removes more Fe per run

Figure 8 shows the cumulative mass of Fe removed during a filtration run and recovered during subsequent backwash of both filters. A total of 47.1 kg Fe were removed from the LOW filter, of which 24.7 kg was recovered in the backwash water (52%). The HIGH filter, operated at a shorter runtime and lower flow rate, removed a total of 17.6 kg Fe from the groundwater. During backwashing, 67% of the Fe (11.7 kg) was recovered, illustrating that the majority of Fe had accumulated in the pores as loose flocs. In both cases, the Fe that is not backwashed results in coating formation, i.e. layers of Fe oxides on the surface of the filter medium. This shift from flocs to coating in the LOW filter seems to explain why the LOW filter did not clog as fast as the HIGH filter, which allowed running it at twice the flow rate (10 m/h vs 5 m/h) and for up to 2-3 times longer. It should be noted that the had a slightly different backwash program, *i*.*e*., lower backwash rate for HIGH, which is indicated by the green markers in Figure 8.

**Figure 8.**
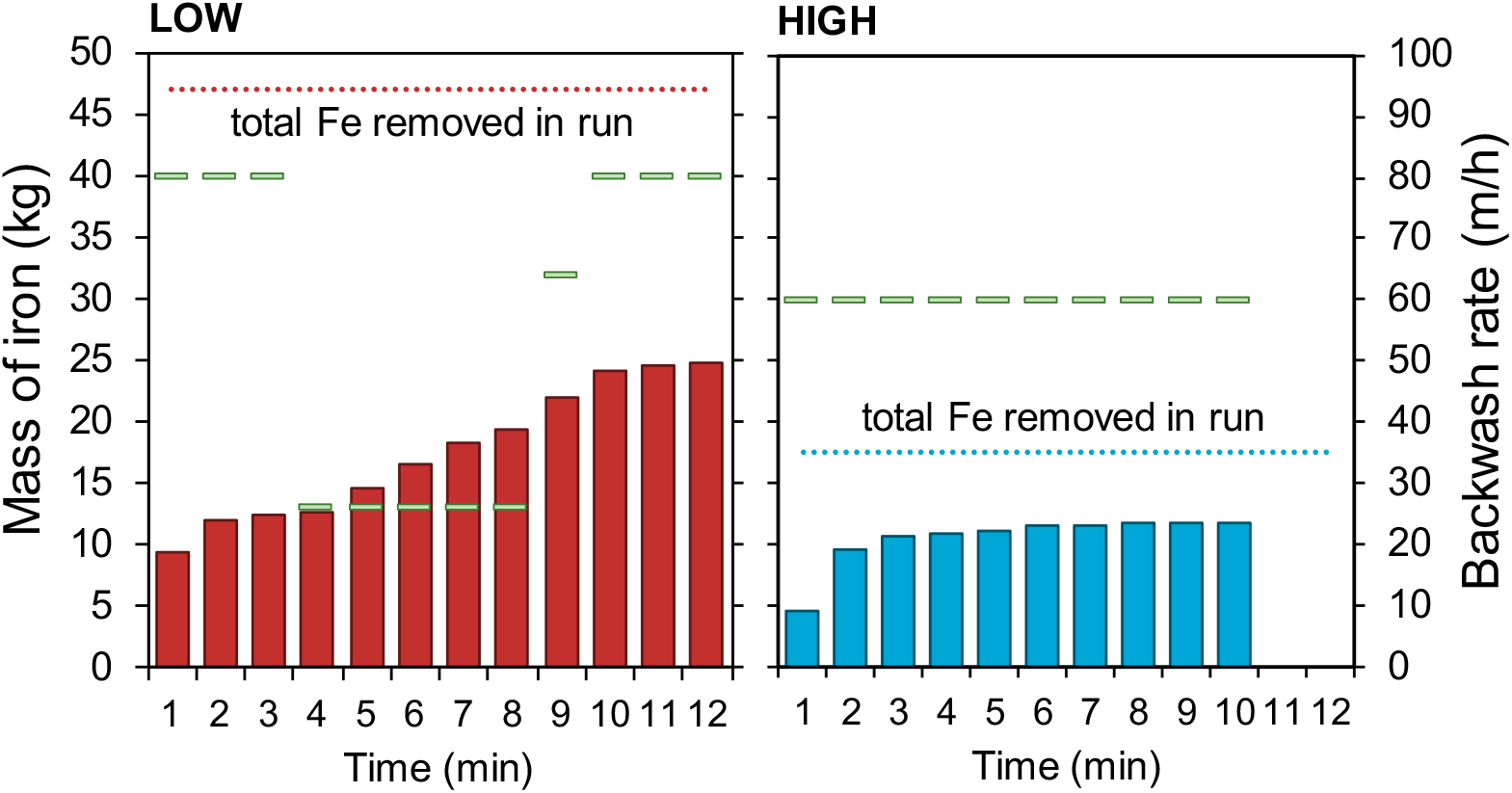
Cumulative mass of Fe over the time of one backwash program (filled bars), backwash rate (green markers) total removed Fe in the run.

### 3.8 No phosphate found in HIGH coating

The composition of Fe solids from the filter bed and backwash water was evaluated with chemical extraction (Figure 9). Fe content was found to be higher in the grain coating of HIGH (50%), compared to LOW (43%). A key difference between the two filter coatings was that while P was abundant in the LOW coating, it was almost absent in the HIGH coating. Instead, P was present in the HIGH backwash solids, which corresponds to the particulate P height profiles in the filter bed. For LOW the reverse was observed, as backwash solids contained lower P content than the coating. Average Ca content was rather stable for all samples (2%) (Figure S2). Si was slightly higher for both coatings than the backwash solids, which might be caused by dissolution of quartz from the sand grains during extraction.

**Figure 9.**
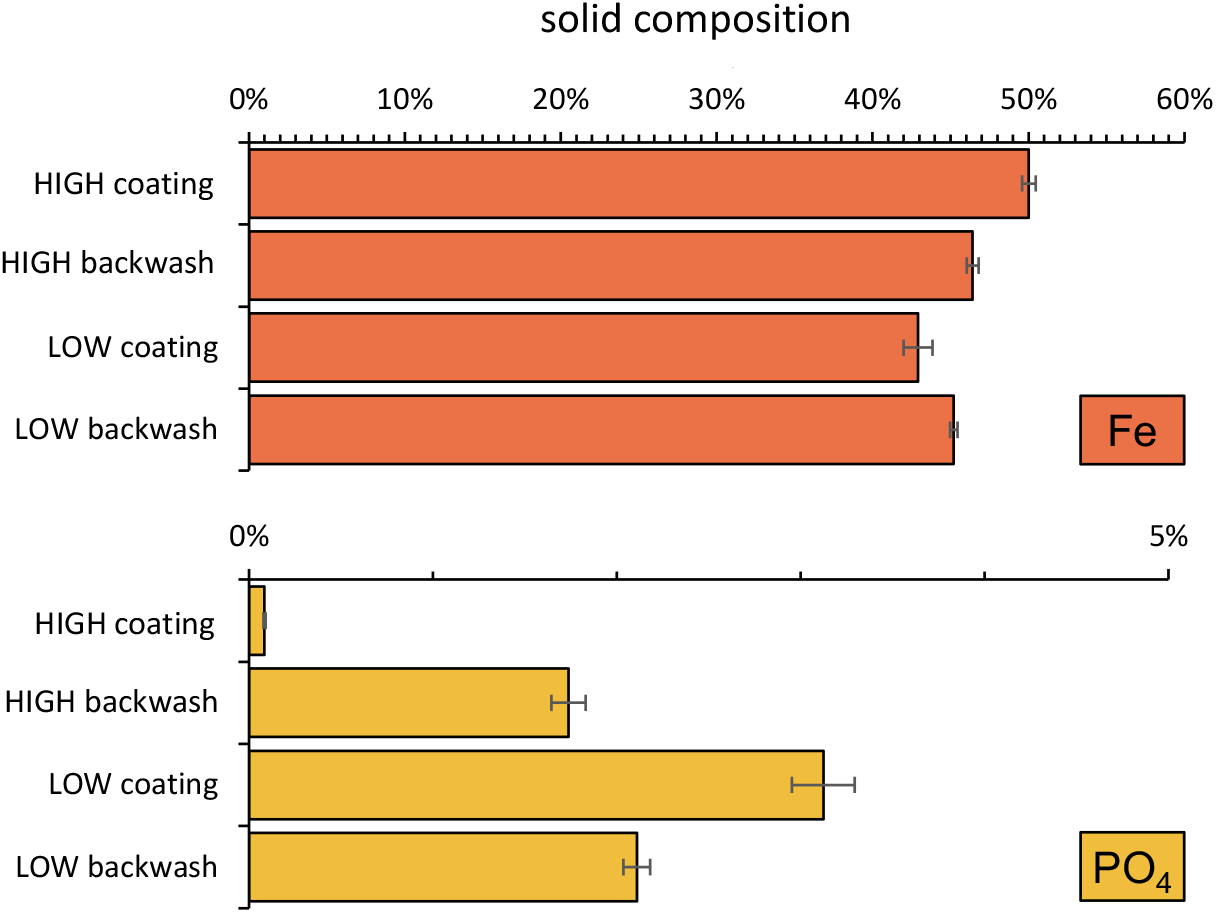
Extractable Fe and P in grain coating and backwash solids of the LOW and HIGH filters. Error bars represent standard deviation of triplicate measurements.

### 3.9 *Gallionella ferruginea* growth in LOW

16s rRNA copies of *G. ferruginea* were detected with qPCR in all samples taken from both filters (Figure 10). The sand coatings obtained from the top and bottom of filter LOW, showed slightly higher copy number than those of the HIGH filter and suggests that *G. ferruginea* was present in both filter coatings. The increase in copy numbers between raw and after gas exchange, illustrates that *G. ferruginea*-containing biofilms grow at the treatment plant, prior to filtration, e.g., in plate aerator, cascades or piping. During sampling, cauliflower-like FeOx structures were observed on the cascades prior to both filters, which could indicate biological activity.

**Figure 10.**
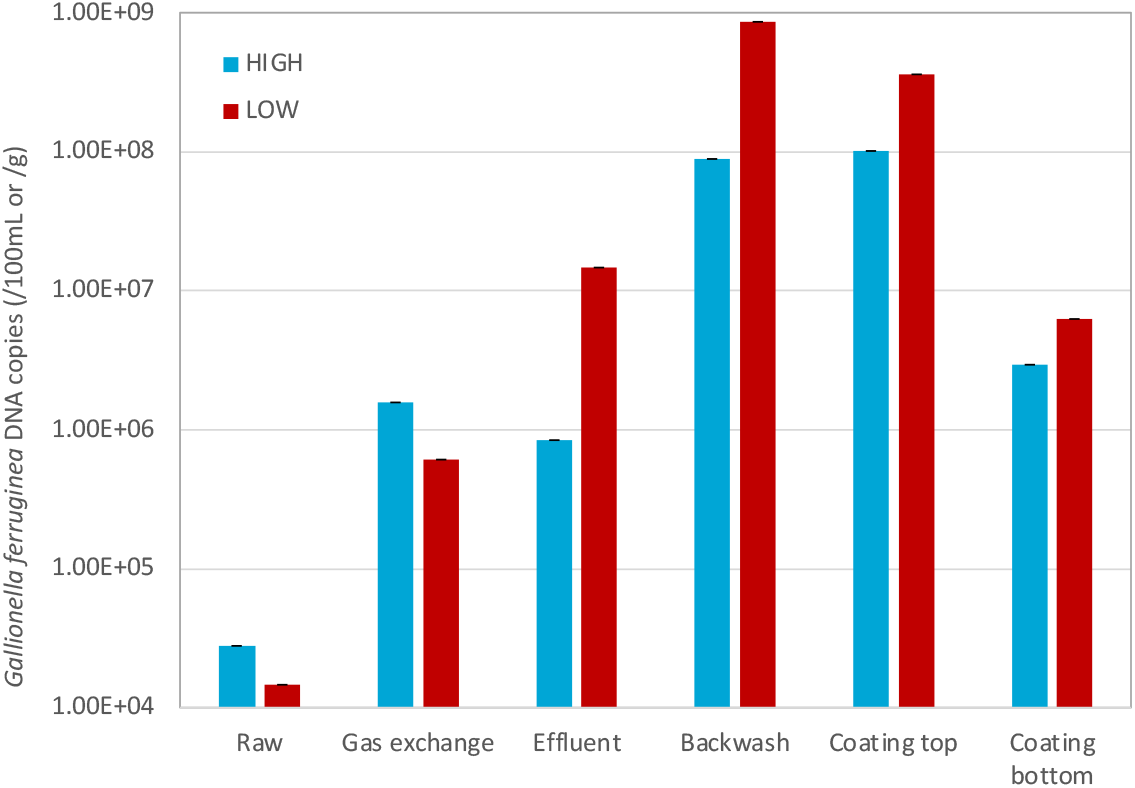
Total number of *G. ferruginea* 16S rRNA copy numbers measured in raw, after gas exchange, effluent, backwash water, and coatings obtained from the top and bottom of the filter.

The effluent of HIGH had a lower copy number than before the filtration (gas exchange). Normalizing per filter run volume (663 m^3^), 4.79*10^12^ copies must have been retained in the HIGH filter (Table 3). In the LOW filter, copies increased after filtration by an order of magnitude, suggesting *G. ferruginea* growth. For the volume produced in that specific run (1800 m^3^), this would correspond to a cumulative wash-out of 2.55*10^14^ copies via the effluent. This is 10 times higher than the influx of *G. ferruginea* DNA copies via the influent (1.10*10^13^), and thus further supporting that the LOW filter harboured biologically active iron oxidation. The absence of wash-out of DNA copies via the effluent of HIGH, suggests that considerable biological growth in this filter is unlikely. However, the backwash samples of HIGH did contain copies in excess to the retained number, making a conclusive mass balance difficult.

**Table 3.**
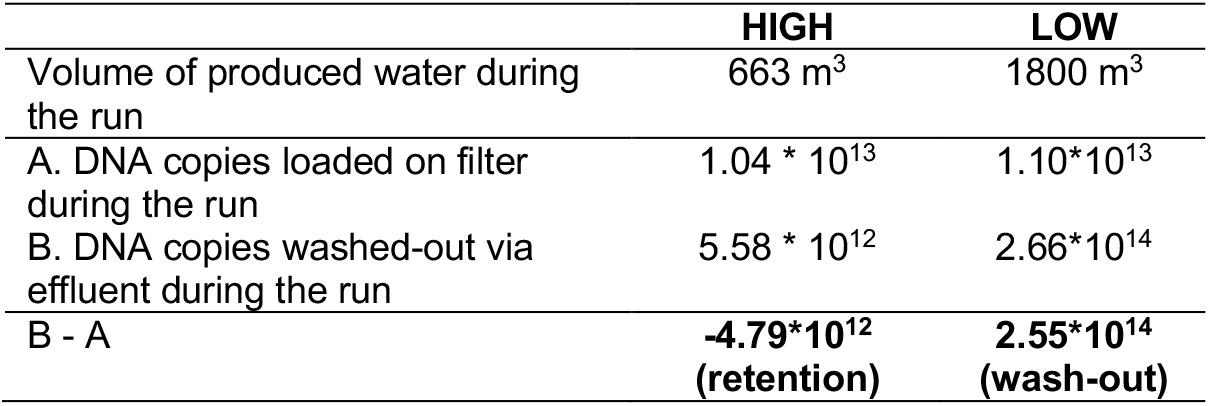
*G. ferruginea 16s rRNA* copy numbers calculated per backwash cycle.

### 3.10. Distinctly different Fe solid morphologies

The differences observed between the Fe removal processes in filters LOW and HIGH were reflected in the morphology of the formed solids. Figure 11 shows digital microscope and ESEM images of the Fe solids on coatings and in backwash water for LOW and HIGH. The filter coating of the LOW filter displayed bright orange coatings with a more porous surface (Figure 11A). The ESEM images (Figure 11C) revealed biogenic stalks (indicator for *G. ferruginea;* Chan et al., 2011) within the coating of LOW, creating large pores, while this was not observed in HIGH (Figure 11D). Thick (>200 µm), dark coatings with smooth surface formed in the HIGH filter (Figure 11B). The backwash solids of filter LOW were dominated by >100 µm long stalks (Figure 11E), which contrasted with the smaller flocs in the HIGH filter’s backwash water (Figure 11F).

**Figure 11.**
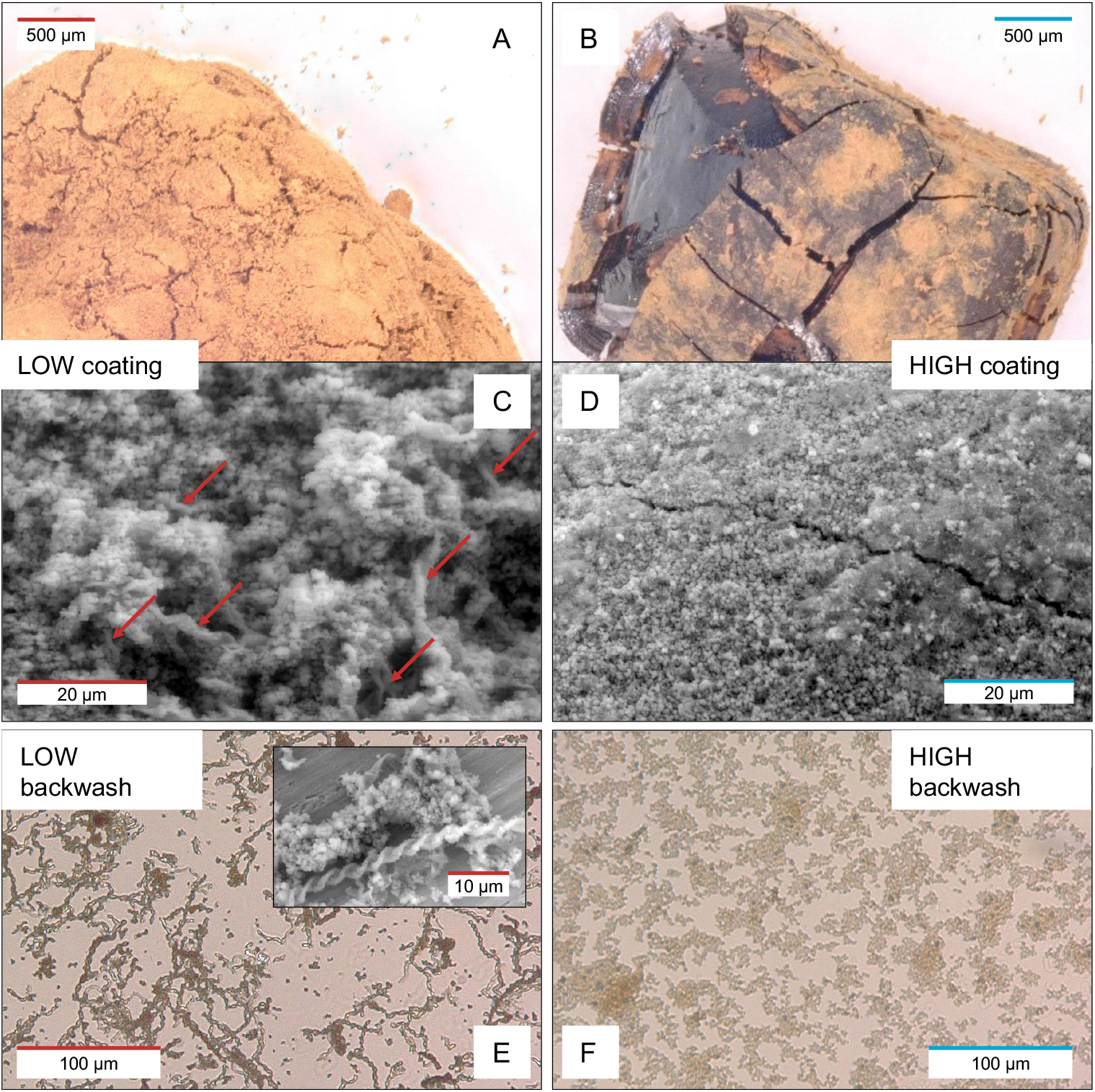
Light microscope and ESEM images filter coatings of LOW (A, C) and HIGH (B, D), as well as back wash solids for LOW (E) and HIGH (F). Arrows in image C indicate incorporation of biogenic twisted stalks in the coating.

The structure of the Fe solids could not be distinguished by XRD as all four samples were poorly crystalline with two broad peaks similar to 2-line-Ferrihydrite (Figure S3). Mössbauer spectroscopy confirmed that all solids are Fe(III) (hydr)oxides of low crystallinity (Figure S4). Fitting of Mössbauer spectra did reveal small differences between the quadrupole splitting of both backwash solids compared to each other and to the coatings, indicating a greater degree of structural disorder in backwash solids in the following order, LOW > HIGH.

### 3.11 Higher adsorption capacity of LOW coating

The rate of heterogeneous oxidation depends on the adsorption of Fe^2+^ to available FeOx. To assess potential difference in adsorption capacity of the FeOx coatings in LOW and HIGH, anaerobic batch experiments were performed in the range of 3-20 mg Fe^2+^/L. Figure 12 depicts the obtained Freundlich adsorption isotherms, with higher K_F_ and n_F_ for LOW (0.18, 1.32) compared to HIGH (0.07, 1.03). Thus, the porous coating on filter grain from filter LOW, as observed with ESEM, provides a more favourable surface for Fe^2+^ adsorption.

**Figure 12.**
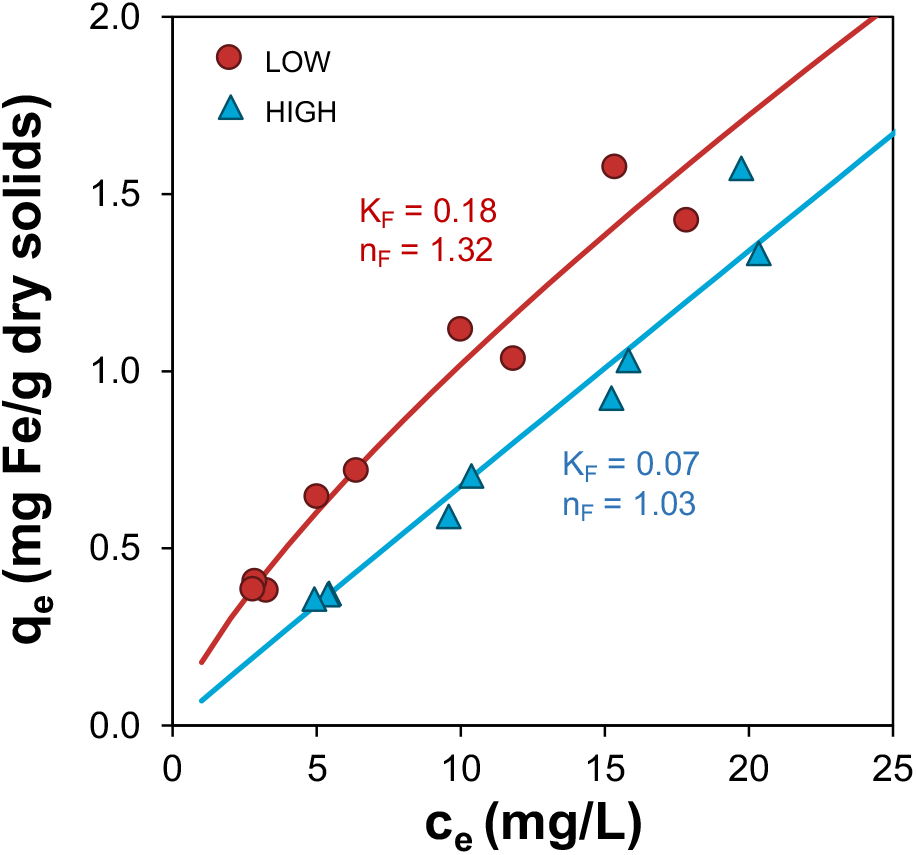
Freundlich isotherm of Fe^2+^ adsorption on filter grains from HIGH and LOW, obtained in anaerobic beaker experiments.

## 4 Discussion

### 4.1 Mass balance reveals fate of Fe

Fe and DO mass balances within one backwash cycle were used to quantify the relative contribution of each oxygen-consuming process and resolve the fate of Fe. The majority of DO in the HIGH filter was used for nitrification, followed by Fe^2+^ oxidation (Figure 13, blue). On the contrary, most of the available DO (5 mg/L) in LOW was used for Fe oxidation, and NH_4+_ oxidation was completely absent. The absence of nitrification could not have been caused by PO_43-_ limitation as seen by others (De Vet et al., 2012), as in the HIGH filter nitrification does onset, albeit P being removed by the particulate Fe prior to filtration. In filter LOW, it seems that the fast consumption of DO by Fe^2+^ oxidation did not allow for the growth of nitrifying organisms in the filter. This preferred oxidation of Fe^2+^ over NH_4+_ is in-line with other studies where Fe and NH_4+_ removals were observed to occur in filters sequentially (Corbera-Rubio et al., 2023). To complete the mass balance in both filters, a minor fraction of DO consumption was attributed to CH_4_ oxidation, assuming complete oxidation to CO_2_.

**Figure 13.**
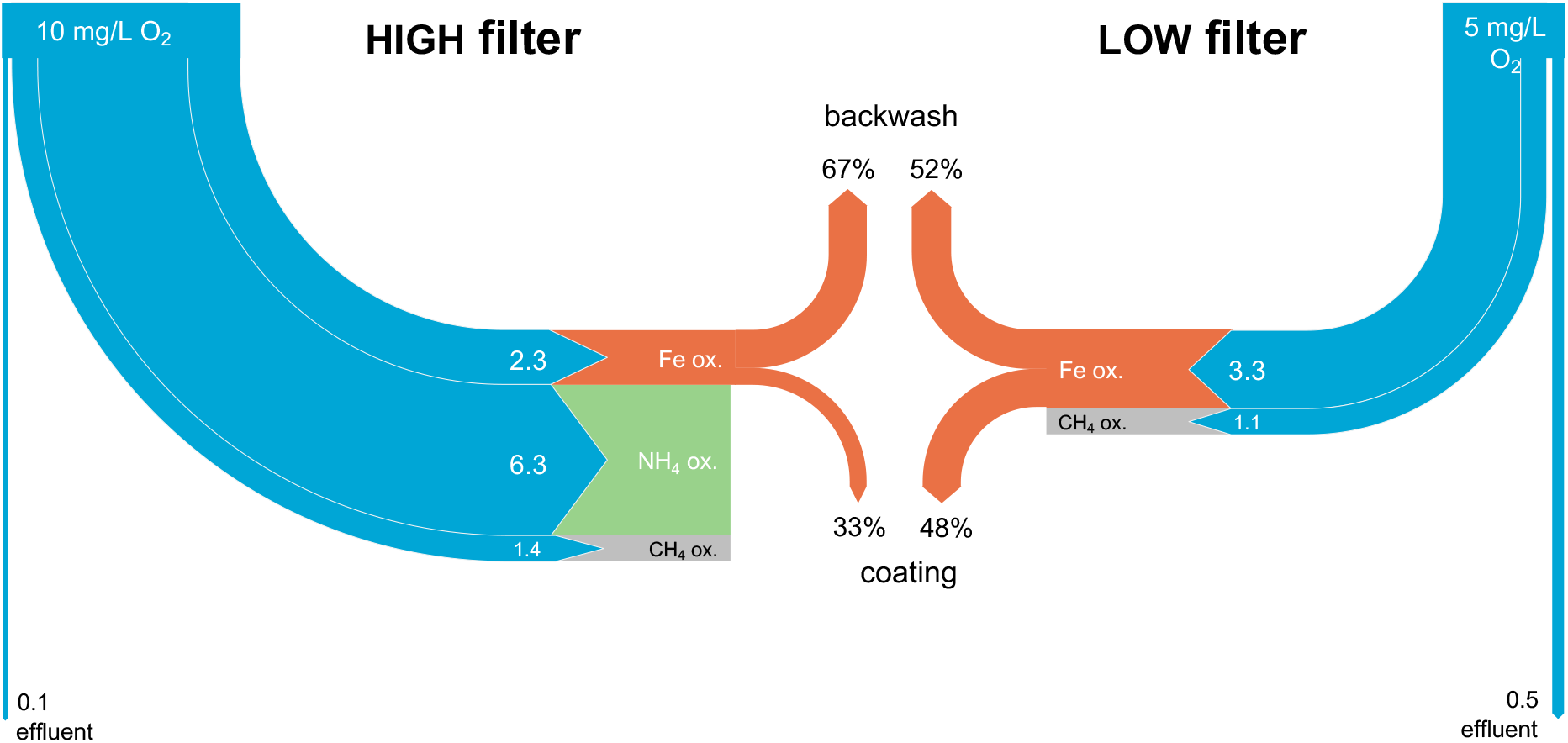
Mass balances of DO (blue) and Fe (orange) in both filters. Concentrations as measured in the supernatant, filter effluent and backwash water.

The mass balance for Fe (Figure 13, orange) shows that 67% of Fe removed during one filter run in HIGH ended up in the backwash suspension, *i*.*e*., the Fe solids were retained in the pore space or loosely attached to the filter medium. FeOx floc formation is associated with homogeneous Fe oxidation, visible as particulate Fe directly after plate aeration and in the supernatant. Flocs are removed in the top meter of the filter, requiring frequent backwashing, and treatment of large volumes of watery Fe sludge. The remaining 34% of the Fe must have contributed to the growth of the filter grain coating. Coating formation in sand filters is usually described as the product of heterogeneous Fe oxidation (Van Beek et al., 2016), where Fe^2+^ is adsorbed on an existing FeOx surface and oxidized by electron transfer to the bulk mineral or surface oxidation (Hiemstra and van Riemsdijk, 2007). Heterogeneous oxidation generates smooth layers of FeOx that grow over time (Haukelidsaeter et al., 2023), in line with the observed morphology in the HIGH filter. FeOx flocs incorporation into the coating is unlikely to have taken place, as these two fractions had different P contents. Also, a significant contribution of biological oxidation to Fe removal in filter HIGH is unlikely, as DNA copies of *G. ferruginea* did not increase after filtration, nor were their signature twisted Fe stalks observed in the Fe solids. We therefore conclude that abiotic oxidation pathways were predominantly responsible for Fe removal in filter HIGH.

In filter LOW, the Fe solids were clearly dominated by the twisted Fe stalks of *G. ferruginea*. These stalks are formed on the surface of the grain but were also observed as the sole Fe solid state in the backwash water. Apparently, the formed Fe stalks partially detached from the surface either during regular operation or backwash. This corresponds to the mass balance (Figure 13), as a large fraction of removed Fe remained as a coating on the filter medium (48%). With the backwash programme in this study, stalks were not completely removed during backwashing and therefore contribute to growth of the visually porous coatings in the LOW filter. It is to further investigate the effect of backwashing on biogenic coating development.

We conclude that biological Fe oxidation and its FeOx products clog pores at a much lower rate than homogeneous oxidation. This agrees with the observed longer runtimes, *i*.*e*., lower backwash frequency, for the LOW filter. Also, due to the accumulation of Fe on the coating, the LOW filter does not produce as much backwash water as the HIGH filter. Instead, periodic extraction of the grains is required to maintain a fixed filter height. These grains are compact and do not need subsequent treatment like the backwash water (*i*.*e*., sedimentation, flocculation, thickening). In addition, FeOx coatings are excellent adsorbents for various reuse applications in drinking or waste water treatment such as PO_43-_ and arsenic adsorption, allowing for their downstream use.

### 4.2 Untangling heterogenous and biological oxidation

In previous studies, heterogenous and biological iron oxidation are often referred to as one mechanism (Mouchet, 1992; Sharma, 2001; Søgaard et al., 2000). This illustrates the challenge of quantifying the contribution of each of these oxidation mechanisms in filters. The presented study has shown that biological Fe oxidation does not play a role in filter HIGH, hence the observed acceleration of Fe oxidation in the filter bed of HIGH is solely caused by heterogenous oxidation (Figure 14A). This twenty-fold jump in rate constant from 0.001 s^-1^ in the supernatant water to 0.02 s^-1^ in the filter bed of HIGH (Figure 14B) can be explained by the fact that heterogeneous oxidation outcompetes homogenous oxidation in the presence of catalytic FeOx surfaces around neutral pH or below (Van Beek et al., 2012). In filter LOW, even higher rates were observed upon entering the filter bed, with a rate constant of 0.045 s^-1^ (Figure 14B). The higher k_LOW,Filter_ than k_HIGH,Filter_, however, contradicts our current understanding of chemical heterogenous Fe_2+_ oxidation, which dictates that higher DO results in higher rates (Tamura, 1980). This suggests that processes that accelerated Fe^2+^ oxidation despite the disadvantageous DO levels dominated in the LOW filter.

**Figure 14.**
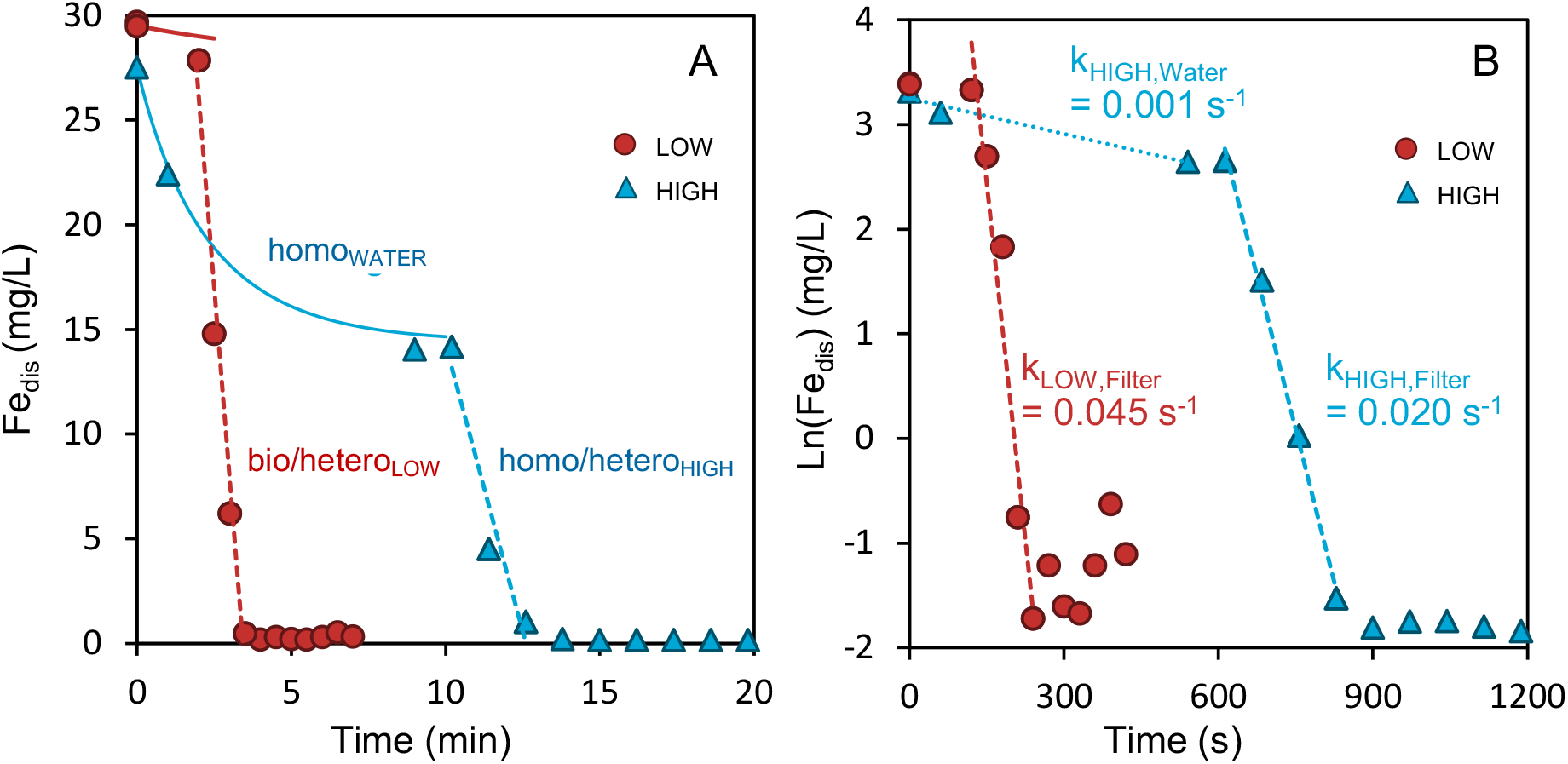
(A) Measured dissolved Fe and (B) natural log of dissolved Fe versus residence time in LOW and HIGH filters. Assuming the maximum residence time prior to filtration of 3 and 10 min for LOW and HIGH, respectively.

ESEM images showed distinctive stalk-like Fe solids, biosignatures of *G. ferruginea* (Chan et al, 2011), in the LOW filter sand coatings as well as in its backwash solids. Also, the LOW filter had 10 times higher DNA copy numbers of *G. ferruginea* in effluent water. This evidence of biological activity points towards the contribution of Fe oxidizing microorganisms in the acceleration of Fe^2+^ oxidation in the filter bed. Fe oxidizing microorganisms are known to contribute to Fe^2+^ oxidation via two different mechanisms, namely (*i*) indirectly due to adsorption on the produced biogenic FeOx coating, or (*ii*) directly via biological oxidation, *i*.*e*., that biology outcompetes chemistry.

Heterogeneous Fe oxidation will inevitably take place in any system with Fe oxidizing bacteria, as biogenic Fe oxides introduce new sorption sites for Fe^2+^, followed by chemical heterogeneous oxidation and precipitation of secondary minerals on the stalk surfaces (Byrne et al., 2018). The specific Fe adsorption capacity (q_e_) was 0.9 mg/kg for the HIGH coating and >2 mg/kg for the biogenic coating in LOW (Figure 12), accounting for the different dissolved Fe concentrations entering the filter beds (Figure 14A). Theoretically, this alone could almost double the heterogenous reaction rate. At the same time, DO concentrations were twice as high in the HIGH filter compared to the LOW filter. According to the established equation of Tamura (1976) for heterogeneous oxidation rate, the lower DO concentration would outbalance the effect of increased FeOx adsorption capacity. In addition, the observed q_e_ was in absence of natural organic matter, suggested to inhibit heterogenous oxidation (de Vet et al., 2011b). Taken together, the indirect pathway of adsorption and heterogenous oxidation on biogenic FeOx solids alone cannot explain the observed acceleration in oxidation.

Therefore, we infer that biological oxidation is responsible for the faster oxidation observed in LOW. Other researchers have also observed that biological oxidation can outcompete heterogeneous oxidation (Druschel et al., 2008; Emerson et al., 2010), although under different circumstances than in our filter (e.g., lower DO). In full-scale groundwater trickling filters, *G. ferruginea* growth was observed previously (de Vet et al., 2011a), which was a different species than found in the source water. Chan and colleagues (2016) showed that Fe stalk formation by *G. ferruginea* does not only offer a mechanism to safely excrete Fe solids preventing cell encrustation, but also enables cell motility. The stalks allow cells to move through the water towards ideal gradients of Fe^2+^ and DO. Applying their conceptual model of mat development on sand filters, a *G. ferruginea* cell on top of a grain would move away from the surface towards the open pore, where Fe^2+^and DO concentrations are highest. We hypothesize that stalk formation might thus be a mechanism to compete with abiotic heterogeneous Fe^2+^ oxidation. Regardless of the precise contribution of heterogenous oxidation, we demonstrate that mild aeration allows to stir filter operation towards biological Fe^2+^ oxidation, and obtain considerably higher oxidation rates than in conventional, intensively aerated filters.

## 5. Conclusions

We operated two parallel full-scale sand filters at different aeration intensities to resolve the relative contribution of homogeneous, heterogeneous and biological Fe removal pathways, and identify their operational controls. The following key conclusions are drawn:

- Lower pH (6.9) and mild aeration (5 mg/L O_2_) enabled Fe^2+^ oxidation at twice the rate compared to the intensively aerated filter (> 10mg/L O_2,_ pH 7.4). Biological Fe^2+^ oxidation dominated in the filter with mild aeration, as supported by 10-fold higher 16s rRNA copies of *G. ferruginea* and ESEM images of their distinctive twisted stalk-like Fe solids.
- Filter clogging was slower for biogenic FeOx than for chemical FeOx, resulting in lower backwash frequencies and yielding four times more water per run.
- Ultimately, our results reveal that biological Fe^2+^ oxidation can be actively controlled and favoured over competing physico-chemical routes. The resulting operational benefits are only starting to be appreciated, with the counterintuitive higher oxidation rates and water yields at lower aeration regimes, and the production of more compact and practically valuable FeOx solids being of outmost interest.

## Acknowledgements

This research is supported by Dunea-Vitens NWO Sand Filtration Programme, grant #17830. Furthermore, we would like to thank Adrie Atsma and Hans Doeve of Vitens, Iulian Dugulan of RID Mössbauer laboratory, Ruud Hendrikx of XRD laboratory and colleagues of the TU Delft Waterlab. DVH and ML were supported by Vidi (#18369) and Veni (#192.252), respectively, of Dutch Research Council NWO.

## Supplementary information

**Figure S1:**
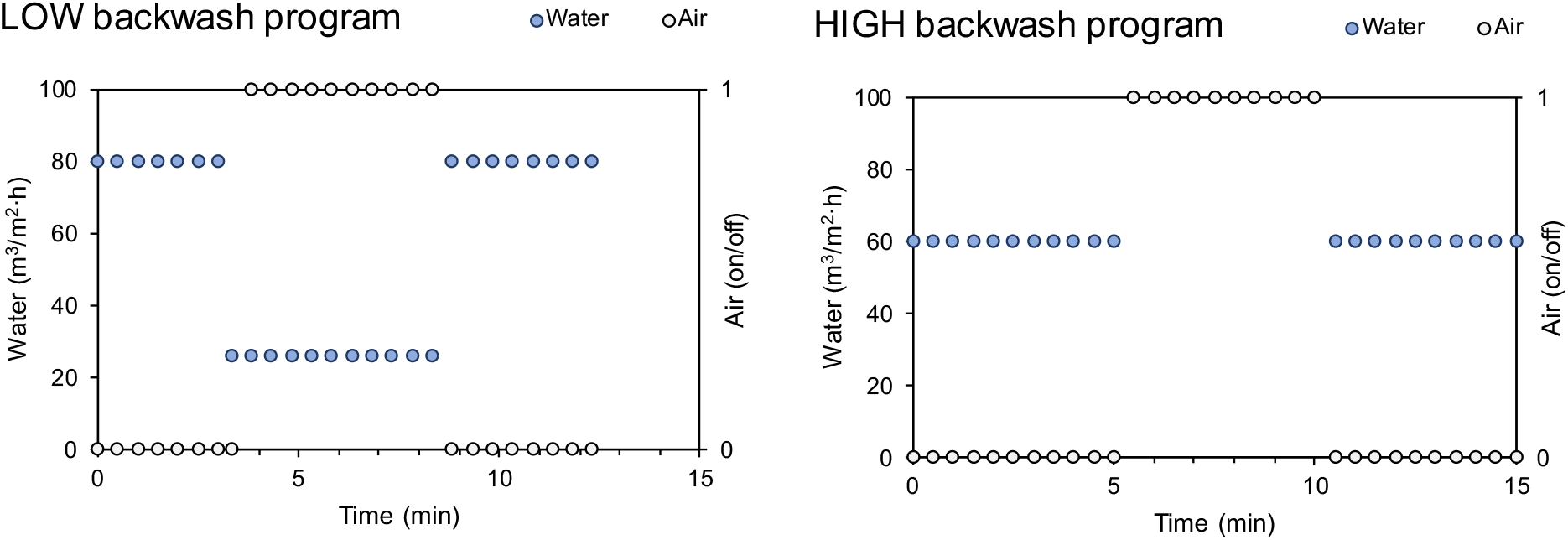
Backwash programs for filter LOW and HIGH

**Figure S2:**
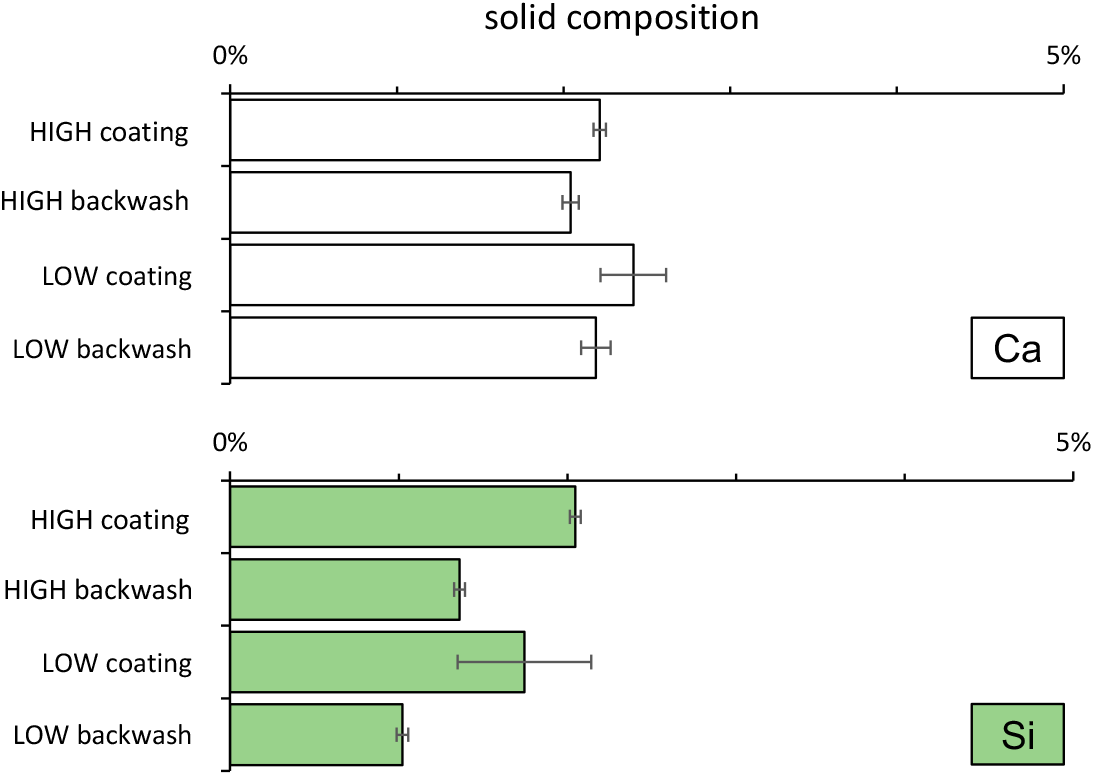
Extractable Ca and Si in grain coating and backwash solids

**Figure S3:**
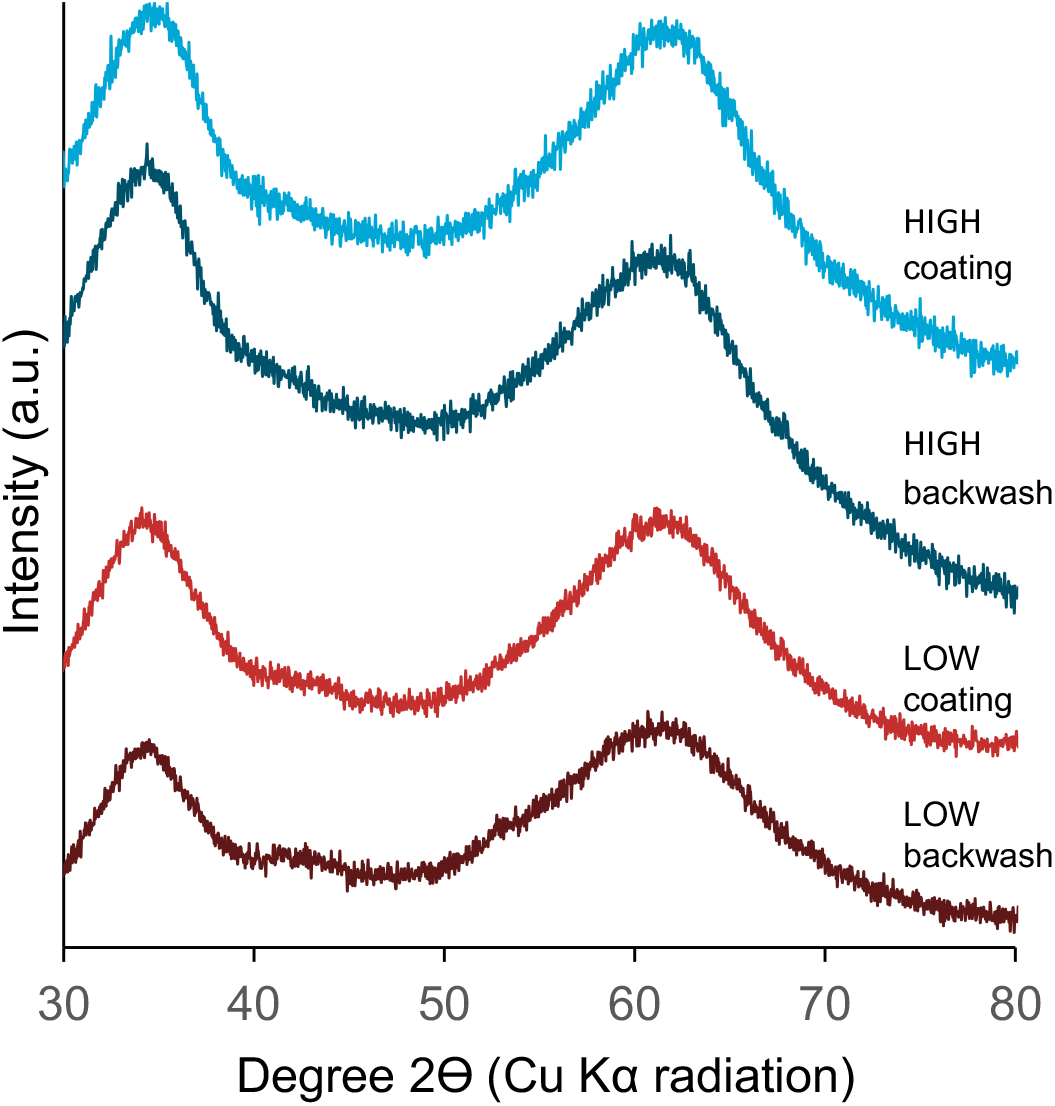
Powder X-Ray Diffractograms

**Figure S4:**
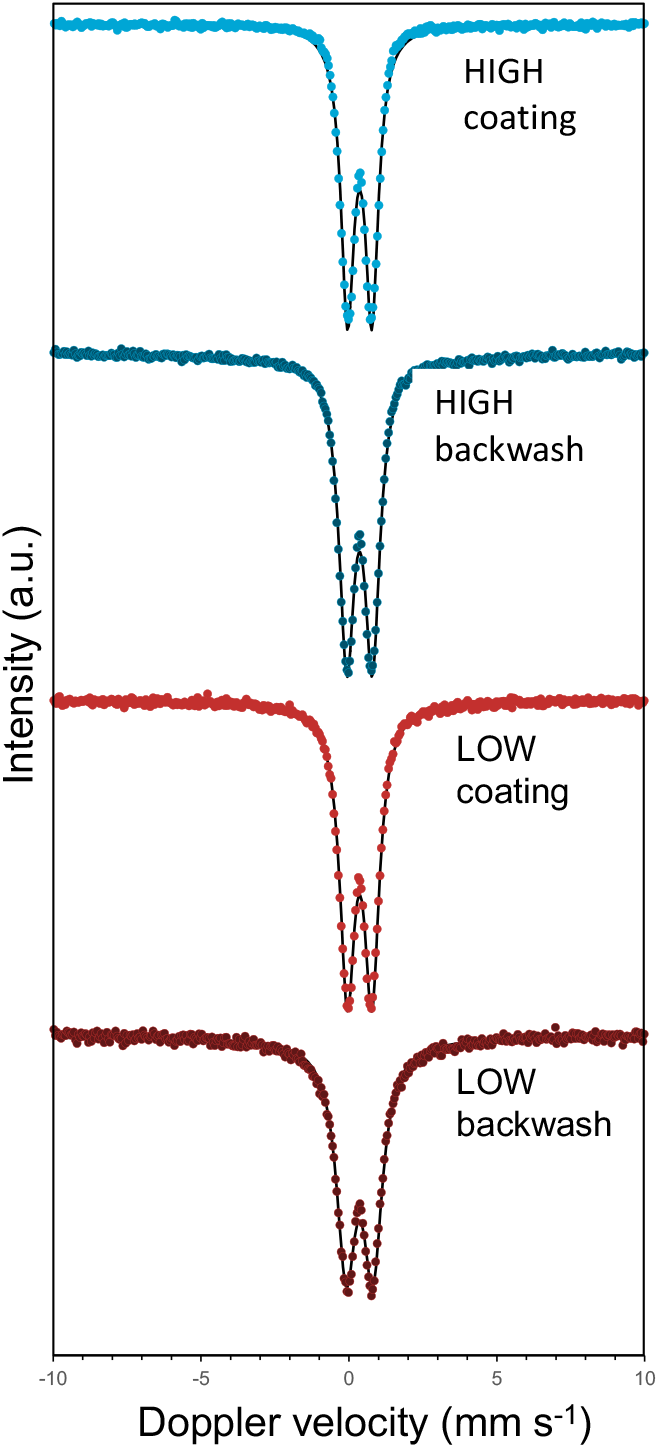
Mössbauer spectra for grain coating and backwash solids from HIGH and LOW

